# Association of Obesity with Health Care Costs: A Mendelian Randomization study

**DOI:** 10.1101/630178

**Authors:** Christoph Kurz, Michael Laxy

**Affiliations:** Institute of Health Economics and Health Care Management, Helmholtz Zentrum München

## Abstract

Causal effect estimates for the association of obesity with health care costs can be biased by reversed causation and omitted variables. In this study, we use genetic variants as instrumental variables to overcome these limitations. We estimate the effect of body mass index (BMI) and waist-to-hip ratio (WHR) on total health care costs using data from a German observational study. We find that the model using genetic instruments identifies additional annual costs of 189€ for a one unit increase in BMI, and additional 1165€ for a 0.1 unit increase in WHR. This is more than two times higher than estimates from linear regression without instrumental variables. We found little evidence of a non-linear relationship between BMI or WHR and health care costs. Our results imply that the use of genetic instruments can be a powerful tool for estimating causal effects in health economic evaluation that might be superior to other types of instruments where there is a strong association with a modifiable risk factor.

## 1 Introduction

Overweight and obesity are global public health concerns in terms of economic costs and effect on health (Wang et al., 2011). The projected global prevalence of obesity will reach 18% in men and surpass 21% in women in 2025 (NCD Risk Factor Collaboration, 2016). Higher body mass index (BMI), expressed in units of kg/m^2^, is a risk factor for type II diabetes mellitus, cardiovascular diseases and certain types of cancer, resulting in shortened life expectancy (Hu et al., 2004). According to the World Health Organization (2000), individuals are classified as obese with a BMI ≥ 30 kg/m^2^ or a waist-to-hip ratio (WHR) ≥ 0.85 for women and ≥ 0.9 for men. While life style factors like physical activity and nutrition are important drivers of obesity, there is also a significant genetic component: at least 34% of BMI variation can be explained by genetic loci (Yang et al., 2011; Maes et al., 1997).

Accurate estimates of health care costs are crucial to judge the trade offs between medical possibilities, their financial viability, as well as quality and fairness in any health care system. Because of practical and ethical issues, many health risk conditions (e.g. obesity) cannot be assessed in randomized controlled trials (RCTs) and cost data collected in RCTs often has limited generalizability (Drummond et al., 2015). On the other hand, observational studies can estimate the relation between health care costs and health conditions, but generally cannot identify causal effects (Cawley and Meyerhoefer, 2012). Many previous studies that estimated the association of obesity with health care costs arise from observational studies (e.g. Finkelstein et al. (2009); Trasande et al. (2009)) that are unable to fully account for unmeasured confounders. For example, factors like social deprivation or bias by self-reported height and weight measures are often not accounted for. In addition, the direction of the omitted variable bias is usually unclear. Cawley and Meyerhoefer (2012) discuss that some people became obese after suffering an injury or chronic depression, and have higher medical costs because of the injury or depression. Carreras-Torres et al. (2018) find that higher BMI increases the risk of being a smoker, but smoking itself lowers BMI (Dare et al., 2015).

The presence of reverse causation and omitted variables motivated the use of instrumental variable (IV) analysis. The aim of the IV method is to exclude a correlation between the explanatory variables and the error term in a regression analysis. This is done by replacing the explanatory variables with other variables that are closely related to them but do not correlate with the error term or are a linear combination of other explanatory variables.

For example, in the case of health care costs of obesity, Cawley and Meyerhoefer (2012) used the BMI of a biological relative as an IV. Compared to a standard ordinary least squares (OLS) model, the authors found a fourfold increase of marginal costs of obesity, and a threefold increase in the marginal costs of one unit of BMI. Studies by Black et al. (2018) and Kinge and Morris (2018) found similar differences between OLS and IV models. They both used the BMI of a biological relative as an IV.

However, a potential concern with this approach is that unobserved characteristics may be correlated with both a person’s own BMI and their relative’s BMI, for example when they live in the same household (Böckerman et al., 2019).

In this paper, we use an IV approach called *Mendelian Randomization* (MR). MR is originally a concept from genetic epidemiology that uses genetic variants as instrumental variables to overcome some limitations in cases where genetic polymorphisms have a well-known effect on modifiable risk factors like BMI (Thomas and Conti, 2004). If the IV assumptions hold, genetic variants can be used to estimate the effect of BMI and WHR on health care costs. Because genotypes are assigned randomly when passed from parents to offspring, the population genotype distribution is assumed to be unrelated to confounders and can therefore serve as valid instruments. This random segregation of alleles, according to Mendel’s law of independent assortment, mimics a natural experiment in which individuals are randomly assigned into groups based on their exposure (Ebrahim and Davey Smith, 2008).

Recent literature suggested that MR can be a powerful tool for economic evaluation (Dixon et al., 2016), however few studies have yet used it (Ding et al., 2009; Fletcher and Lehrer, 2011; Norton and Han, 2008; Cawley et al., 2011; Willage, 2017; von Hinke et al., 2016; Böckerman et al., 2019).

In this study we use Mendelian Randomization to estimate the effect of BMI and WHR on total annual health care costs and compare the results to a simple OLS model.

The rest of the paper is organized as follows. In Section 2 we first outline our data in Subsection 2.1. We then detail the MR approach in Subsection 2.2 and potential problems regarding the validity of this approach in Subsections 2.3 to 2.5. We present results in Section 3. Finally, we conclude in Section 4 with implications of the results.

## 2 Material and Methods

### 2.1 Data

We use data from the KORA (Cooperative Health Research in the Augsburg Region) F4 study (2006-2010, n=3080). KORA is a population-based research platform in the region of Augsburg, a city in the south-west of Germany, and two surrounding districts running population-based epidemiological studies. A more detailed description of KORA can be found in Holle et al. (2005). Participants were interviewed at the study center regarding demographic and disease-related parameters, health care utilization and medications. Weight and height measurements were performed by trained staff. Education was classified in 3 groups: *basic education* (≤ 9 years of schooling), *medium education*, and *higher education* (≥ 12 years of schooling, required to enter university). Alcohol consumption was measured as average daily dose (in grams), physical activity was categorized in *high* (>2h/ week), *moderate* (up to 2 hours/week), and *no activity*. The groups of working status were: *working, unemployed, retired*, and *other*.

Resource utilization regarding physician visits, in- and out-patient hospital treatments, rehabilitations and medication were assessed by extrapolating the utilization of the asked time horizon (7 days – 12 months) to one full year. Health care costs were estimated by multiplying utilization of physician visits, in- and out-patient hospital treatments, and rehabilitations with the unit costs provided by Bock et al. (2015). The costs of prescription medication was estimated using pharmacy retail prices, based on patient information on medication name and national drug codes. All costs were adjusted to 2011 prices in €. Table 1 presents an overview of the data set. Underweight individuals (BMI < 18.5, *n* = 10) were removed because this group is especially vulnerable to bias due to other pre-existing disease (Kent et al., 2017). Observations with missing or implausible information were also removed, resulting in a sample size of *n* = 2779.

**Table 1:** Summary statistics for the KORA F4 data set. Health care costs are expressed in 2011-€.

|  | Overall |
| --- | --- |
| Annual total health care costs (mean (sd)) | 1,559 (3,645) |
| BMI (mean (sd)) | 27.7 (4.8) |
| WHR (mean (sd)) | 0.88 (0.9) |
| Age (mean (sd)) | 56.2 (13.2) |
| Sex |  |
| Female (n (%)) | 1,410 (50.7) |
| Male (n (%)) | 1,369 (49.3) |
| Alcohol consumption g/day (mean (sd)) | 14.42 (19.5) |
| Education |  |
| basic (n (%)) | 1,447 (52) |
| medium (n (%)) | 675 (24) |
| higher (n (%)) | 657 (24) |
| Working status |  |
| working (n (%)) | 1,829 (66) |
| unemployed (n (%)) | 55 (2) |
| retired (n (%)) | 574 (21) |
| other (n (%)) | 321 (12) |
| Physical activity |  |
| not active (n (%)) | 1,829 (66) |
| moderate (n (%)) | 55 (2) |
| active (n (%)) | 574 (21) |
| Observations n | 2,779 |

Single nucleotide polymorhpisms (SNPs) were available from Affymetrix Axiom genotyping for all participants. These genotypes were entered into imputation via the Michigan imputation server with HRC reference panel version r1.1 (McCarthy et al., 2016). We selected the 97 genetic loci that are highly associated (*p* < 5 × 10^*-*8^) with BMI variation, based on a recent consortium-based genome-wide association study (GWAS) by Locke et al. (2015). For WHR, we used the 48 SNPs found by the GWAS study by Shungin et al. (2015). Some SNPs are relevant for both BMI and WHR. Gene selection based on large GWAS make ideal candidates for MR studies (Taylor et al., 2014). All analyses were conducted in R (R Core Team, 2019; Kleiber and Zeileis, 2008).

### 2.2 (One sample) Mendelian randomization and two-stage least-squares

Mendelian randomization is a type of IV analysis that uses genetic variants as instruments. The random allocation of genetic variants at conception mean that these variants are less likely to violate some of the assumptions of IV analysis than non-genetic instruments, even describing it as nature’s randomized controlled trial (Smith, 2006).

To estimate the impact of BMI and WHR on medical spending, we used a two-stage least-squares (2SLS) model. In the first step of the 2SLS approach, the endogenous regressor of interest (BMI or WHR) is regressed on all valid instruments. Since the instruments are exogenous, this approximation of the endogenous variables will not correlate with the error term.

In the second step, the outcome (total annual health care costs) is regressed on the fitted values of obesity from the first stage.

Stage 1: Regress independent variable *X* (BMI or WHR) on instruments *Z* (genetic loci):

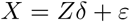

Stage 2: Regress *Y* (medical expenditures) on the predictions 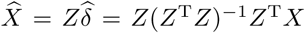 from the first stage:

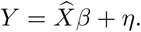

The 2SLS estimates are then

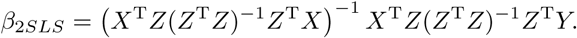

All 2SLS models control for age, sex, alcohol intake, education, working status, and physical activity.

Our approach, using individual level data, is called *one-sample* MR. Another popular approach is two-sample MR, using summarized data from GWAS consortia in a meta analysis. The main advantage of two-sample MR is increased statistical power, but the quality of the pooled results is dependent on that of the individual studies. That means, two-sample MR has to use the summary results presented, even when they have been adjusted for covariates that you would rather they had not been adjusted for. It is important to not view genetic susceptibility for complex conditions in isolation but along with lifestyle and environmental factors (Sugrue and Desikan, 2019), a major advantage of one-sample MR. See Lawlor (2016) for a detailed comparison of one-sample and two-sample MR.

For comparison, we compute linear OLS non-IV models that regress health care costs on BMI and WHR and again adjust for the same covariates as before. We use the Durbin-Wu-Hausman endogeneity test (Hausman, 1978) to evaluate whether there is any evidence that the instrumental variable estimate differs from the OLS estimate. In this test, a significant results indicates disagreement between OLS and IV estimates.

### 2.3 Pleiotropy

A common problem in MR analyses is the presence of pleiotropic effects, i.e. a genetic variant has associations with more than one risk factor on different causal pathways (Lawlor et al., 2008). This can be especially problematic for polygenic risk factors like BMI, where the influence of genetic variants is less specific (VanderWeele et al., 2014). To test violations of the IV assumptions, we used three approaches. First, we checked, in the Phenoscanner database (Staley et al., 2016), if genes related to BMI and WHR are also significantly associated with other determinants of health care costs. We found that some SNPs are related to obesity-related illnesses like type 2 diabetes and high blood pressure. As in Böckerman et al. (2019) we assume that the associations with obesity-related conditions occur because of the SNPs’ association with high BMI and WHR, but we cannot definitely rule out other pathways. In a second check, we used the MR-Egger method (Bowden et al., 2015) to test for pleiotropy. The MR-Egger method performs a regression of the SNP-outcome associations on the SNP-exposure associations with the intercept left unconstrained to test for evidence of bias-generating pleiotropy. As a third test for pleiotropy we conducted Sargan’s test for overidentification (Sargan, 1958). The null hypothesis in this test is that all exogenous instruments are in fact exogenous and uncorrelated with the model residuals, and all over-identifying restrictions are therefore valid.

### 2.4 Invalid instruments

As the number of biomarker-associated variants is constantly increasing through GWAS, selection of the most appropriate instruments is an important issue (Swerdlow et al., 2016). Using too many genetic variants as instruments can lead to spurious estimates and increased type 1 error rates. This implies that, even if a set of multiple instruments is valid, (i.e. they are not associated with confounding factors, have no direct effect on the outcome, and are at least weakly associated with the exposure) the 2SLS estimator can still be biased towards the conventional regression estimate (Davies et al., 2015; Bound et al., 1995). A weak instrument only explains a small proportion of the variation in the risk factor. Using many weak instruments can still result in weak instrument bias (Bound et al., 1995; Staiger and Stock, 1997).

For this, Belloni et al. (2012) suggest selecting optimal instruments in the first stage regression by least absolute shrinkage and selection operator (LASSO) variable selection. A similar approach with explicit application to MR was propsed by Kang et al. (2016). Additional work by Windmeijer et al. (2018) recommends to use the adaptive LASSO to retain the oracle properties (Zou, 2006). In the adaptive LASSO for the first stage, the goal is to minimize

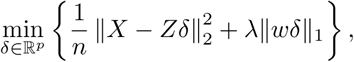

where *λ* is a penalization parameter obtained by cross-validation, and 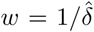 is an initial estimate of *δ* (e.g., obtained by least squares). The LASSO procedure shrinks many *δ* parameters to effectively zero, pruning them out of the regression. According to Keele and Morgan (2016) we additionally perform an F-test on the first stage regression to test for weak instruments.

### 2.5 Non-linearity

Previous research found evidence of a non-linear relationship between BMI and health care costs (Laxy et al., 2017; Cawley and Meyerhoefer, 2012), and MR is often unable to detect such non-linearities because genetic variants usually only explain a small percentage of the variance in the risk factor (Staley and Burgess, 2017). To test for non-linear effects of BMI and WHR on health care costs in our sample, we fit a generalized additive model (GAM) (Hastie and Tibshirani, 1986) where we regress the linear predictor (health care costs) linearly on unknown smooth functions of some predictor variable (BMI or WHR), adjusted for the same covariates as in the MR model.

## 3 Results

LASSO selects 64 valid genetic instruments for BMI and 15 for WHR. The relevant SNPs and their first stage regression estimates are shown in Figure 1. In both IV analyses, we find little evidence of weak instruments, because the weak instruments test *p*-value is < 0.0001 in both cases and the F-statistics are above the traditional threshold of 10 (Stock and Yogo, 2005).

**Figure 1:**
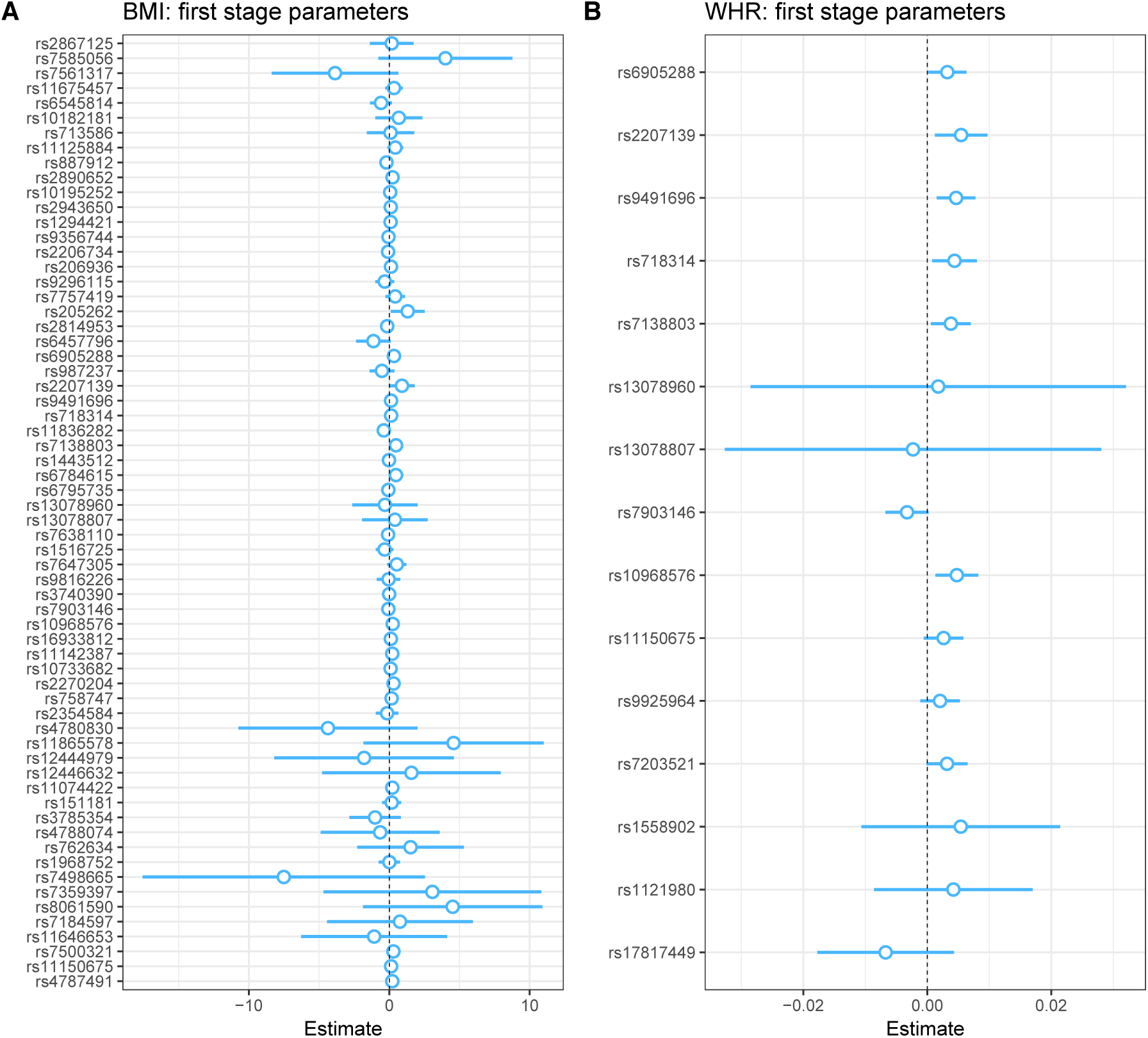
Forestplot for the parameter estimates and 95% confidence intervals of the first stage regression for (A) BMI and (B) WHR. Only genetic variables (SNPs) are shown.

The Sargan test does not provide evidence of pleiotropic effects in both BMI and WHR IV models (see *p*-values in Table 2 and Table 3). This is supported by the MR Egger intercept test (see Figure 2). The Durbin-Wu-Hausman test, however, suggests no strong evidence of differences between the OLS and IV estimates.

**Table 2:** The effect of BMI on annual total health care costs using an OLS model (left column) and an IV model with genetic variants (right column). The IV results are from the second stage of the 2SLS approach. Standard errors in parenthesis.

| Dependent variable: | *OLS* | *IV* |
| --- | --- | --- |
|  | Annual total health care costs | Annual total health care costs |
| Age | 39.2***<br>(5.8) | 32.6***<br>(7.6) |
| Sex = Female | 188.5<br>(149.9) | 255.5<br>(159.2) |
| BMI | 75.5***<br>(14.9) | 188.5**<br>(84.3) |
| Education |  |  |
| basic | Ref. | Ref. |
| medium | −19.2<br>(171.9) | 108.9<br>(197.5) |
| higher | −400.7**<br>(178.9) | −205.2<br>(230.8) |
| Alcohol consumption | −1.2<br>(3.7) | −1.3<br>(3.7) |
| Working status |  |  |
| working | Ref. | Ref. |
| unemployed | 368.6<br>(488.5) | 303.2<br>(495.9) |
| retired | 187.5<br>(182.8) | 130.5<br>(189.4) |
| other | 47.2<br>(225.4) | −4.2<br>(230.8) |
| Physical activity |  |  |
| not active | Ref. | Ref. |
| moderate | −59.4<br>(160.4) | 29.7<br>(174.8) |
| active | −116.6<br>(185.6) | 61.8<br>(228.7) |
| Constant | −2,895.6***<br>(586.6) | −5,894.1***<br>(2,279.4) |
| Observations | 2,779 | 2,779 |
| Weak instruments test *p*-value |  | <0.0001 |
| First stage F-statistic |  | 12.9 |
| Durbin-Wu-Hausman test *p*-value |  | 0.17 |
| Sargan test *p*-value |  | 0.67 |
| *Note:* * *p* <0.1; ** *p* <0.05; *** *p* <0.01 |  |  |

**Table 3:** The effect of WHR on annual total health care costs using an OLS model (left column) and an IV model with genetic variants (right column). The IV results are from the second stage of the 2SLS approach. Standard errors in parenthesis.

| Dependent variable: | *OLS* | *IV* |
| --- | --- | --- |
|  | Annual total health care costs | Annual total health care costs |
| Age | 34.1***<br>(6.1) | 20.9<br>(16.7) |
| Sex = Female | 684.3***<br>(195.6) | 1,436.3<br>(904.9) |
| WHR | 4,872.1***<br>(1,133.2) | 11,649.9<br>(8,042.3) |
| Education |  |  |
| basic | Ref. | Ref. |
| medium | −48.5<br>(171.8) | 29.8<br>(195.8) |
| higher | −431.2**<br>(178.8) | −292.2<br>(243.0) |
| Alcohol consumption | −2.2<br>(3.7) | −3.6<br>(4.1) |
| Working status |  |  |
| working | Ref. | Ref. |
| unemployed | 313.5<br>(489.6) | 176.0<br>(518.5) |
| retired | 176.9<br>(183.2) | 109.2<br>(200.7) |
| other | 13.5<br>(226.1) | −80.9<br>(253.2) |
| Physical activity |  |  |
| not active | Ref. | Ref. |
| moderate | −74.2<br>(160.5) | −12.1<br>(177.3) |
| active | −134.0<br>(185.8) | 7.4<br>(250.1) |
| Constant | −5,517.9***<br>(586.6) | −11,949.9**<br>(2,279.4) |
| Observations | 2,779 | 2,779 |
| Weak instruments test *p*-value |  | <0.0001 |
| First stage F-statistic |  | 12.0 |
| Durbin-Wu-Hausman test *p*-value |  | 0.39 |
| Sargan test *p*-value |  | 0.97 |

*Note:* \**p*<0.1; \*\**p*<0.05; \*\*\**p*<0.01

**Figure 2:**
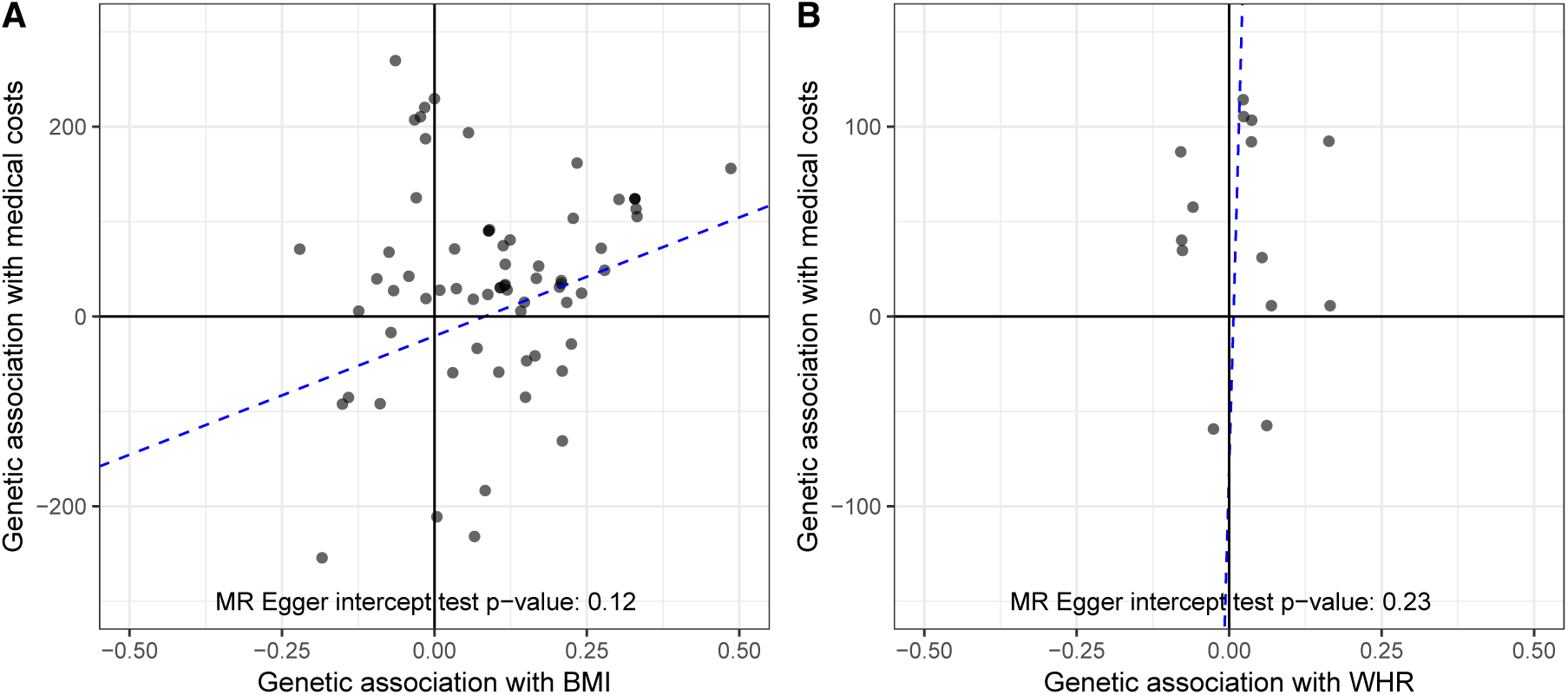
Plot of the gene-outcome vs gene-exposure regression coefficients for (A) BMI and (B) WHR. The dashed line represents the MR Egger regression estimate. Some outliers are not shown.

Table 2 presents the results of the association between BMI and health care costs and Table 3 the results of WHR and health care costs. In the case of BMI, the OLS model finds an effect of 75.5€ for a one unit increase of BMI. In contrast, the IV model estimates an effect of 188.5€ for a one unit increase of BMI.

For WHR, a 0.1 increase of WHR is associated with increased costs of 487€ for the OLS model and 1165€ for the IV model. In both cases, the IV model using genetic instruments estimates the effect on costs almost 2.5 times higher than the OLS model. Other significant cost drivers or savers are age and higher education for the BMI model, and, in addition, sex in the WHR model. Especially in the WHR case (see Table 3), many associations lose its significance at the 0.05 level when moving from OLS to IV analysis.

We also detect no proof of strong non-linear effects of BMI or WHR on health care costs in our sample, according to the GAM models, see Figure 3.

**Figure 3:**
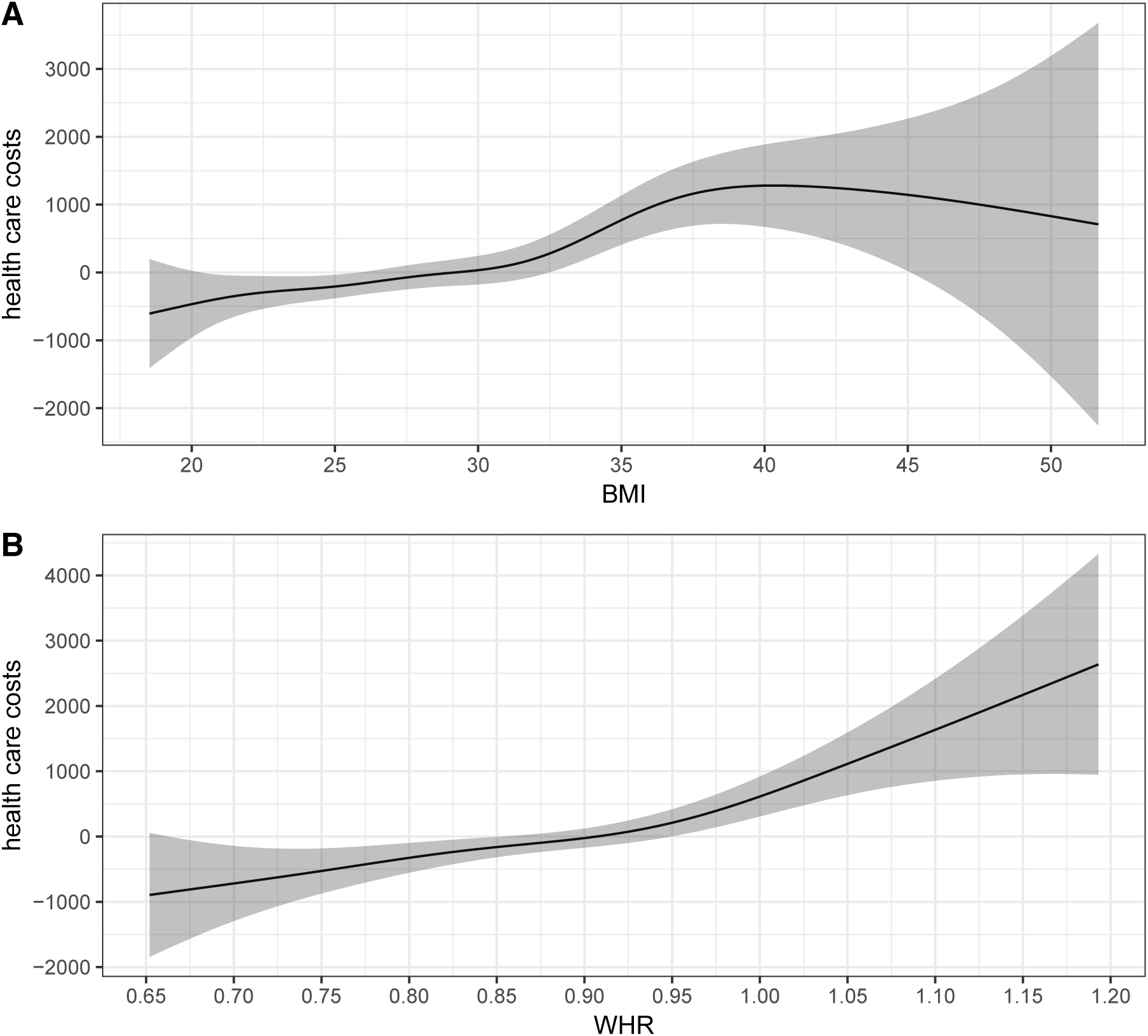
Plots of the smooth function from the GAM for (A) BMI and (B) WHR. The shaded area represents the 95% confidence band.

## 4 Discussion

This paper is one of the first that uses a one-sample MR approach to estimate the marginal health care costs of a prevalent clinical condition. MR offers new opportunities for reliable causal inference in health economic research within the framework of observational research designs. Using MR, our findings indicate that a one unit increase of BMI increases total medical spending by 188.5€ and a 0.1 unit increase of WHR increased total medical spending by 1165€. The OLS model, that doesn’t use instrumental variables, only finds additional spending of 75.5€ for BMI and 487€ for WHR. This demonstrates that a MR study with genetic instruments may detect more hidden bias than a non-IV analysis, leading to higher estimated effects. It is important to note that OLS estimates average treatment effects, while IV estimates local average treatment effects. This may partially account for the differences in the effect estimates (Imbens and Angrist, 1994). Our study confirms the results in Cawley and Meyerhoefer (2012), Black et al. (2018), and Kinge and Morris (2018) that imply much higher costs of obesity in the IV analysis compared with the non-IV analysis. These studies use the BMI of the respondent’s oldest child as instrument and report that the IV-estimate for the association between BMI and health care costs is between two to four times higher than the non-IV approach. One possible explanation for the even greater difference between the IV and non-IV estimates in the US by the Cawley and Meyerhoefer (2012) study might be that subgroups with higher obesity prevalence have reduced access to health care in the US, while Germany has a social insurance system covering almost all citizens (Davis and Ballreich, 2014). However, as it is likely that, due to the shared environmental exposures, a child’s BMI is correlated with relevant covariates (Dubois et al., 2012), the use of such non or quasi genetic IVs is discussed controversial and might lead to biased estimates as well.

Another advantage of our data is that height, weight, waist and hip measurements of individuals in the sample were performed by trained staff. This is more accurate than self-reported measurements where people tend to under-estimate their weight and over-estimate their height, resulting in an under-estimation in BMI (Gorber et al., 2007).

Because participants in our study were asked to self-report their health service usage over the past 3–12 months, recall bias cannot be excluded. Health care utilization was priced with average reference value, therefore actual costs might deviate. However, this should not influence the validity of the study results (Evans and Crawford, 1999). In addition, we were unable to consider cost categories such as presenteeism, premature death or out-of-pocket payments for medication. Although the problem of recall bias and average unit costs are likely to lead to an underestimation of absolute costs, this should not strongly influence the cost increase between BMI and WHR units that were the focus of this study.

Other studies found similar results. Within the BMI range of 25 to 45 kg/m^2^, a study in the United States by Wang et al. (2006) described an increase in medical and pharmaceutical costs of $202.3 per BMI unit. A review of 75 international studies by Kent et al. (2017) reported a median increase in mean total annual health care costs of 36% for obese individuals compared with individuals of healthy weight. All of these studies rely on traditional non-IV estimation.

Having weak or invalid instruments can be a problem in MR analysis. Davies et al. (2015) show that 2SLS is especially vulnerable to weak instrument bias in MR and propose the limited information maximum likelihood and the continuously updating estimator as unbiased alternatives to 2SLS. However, these methods are difficult to implement and interpret. For this reason, we used the adaptive LASSO variable selection in the first state of 2SLS to rule out invalid and weak instruments. The SNPs we used have been discovered in GWAS and should be valid.

The Durbin-Wu-Hausman test *p*-value suggests no significant difference between OLS and IV estimates. However, the effects being estimated by the two methods may not be the same–the MR estimate reflects the effects of lifelong perturbations in the risk factor, whereas OLS regression results may reflect more acute effects (Davies et al., 2018). According to Burgess and Thompson (2015), it would be fallacious to assume that a non-significant result means the OLS estimate is unconfounded. MR estimates are almost always less precise and have wider confidence intervals than OLS regression, so tests for difference often have low statistical power.

It must also be noted that the KORA F4 study was part of the GWAS consortium that discovered the relevant SNPs for BMI in the study by Locke et al. (2015). Because the same data was used at the GWAS discovery stage and in our analysis, it is possible that chance correlation between SNPs and confounders can lead to overestimation of the SNP-trait effect (Haycock et al., 2016). This is the so-called winner’s curse or Beavis effect (Göring et al., 2001). However, the KORA F4 study was only a small fraction of the more than 300,000 individuals in this GWAS, so it should not strongly affect the power of our analysis.

There are alternative ways to include genetic information into the IV model. For example, an allele score can be generated from the genetic variants and included as a single instrument in the association model. This allele score is calculated per individual as the weighted or unweighted sum of the number of risk or trait increasing alleles (Teumer, 2018). However, Palmer et al. (2012) showed that an unweighted score has lower power than adding multiple IVs into the 2SLS, and using an appropriately weighted allele score performs similarly to adding each valid SNP as an instrument. The effect estimate is a little less biased when using a weighted allele score but has less precision (and power) compared to the multiple IV 2SLS estimator that we used.

Burgess et al. (2014) note that instrumental variable estimates using a linear model may not reflect causal effects for large changes in the exposure. In our case we find the relationship between BMI and costs and WHR and costs approximately linear and therefore the linear model is justified.

In the 2SLS estimator, we assume normal distribution of cost data which is not present because of skewness, positivity, and heavy tails. This can also affect the validity of the standard errors and confidence intervals (Guo et al., 2018; Kang et al., 2018). Still, the use of a linear model is justified because the skewness and tail distribution are not extreme and the large sample size guarantees near-normality of sample means because of the Central Limit Theorem (Mihaylova et al., 2011). However, more sophisticated generalized linear models using different error distributions could be examined (Duan et al., 1983; Kurz, 2017).

In conclusion, we showed that MR can be a viable tool in health economic analyses. We found more than two times higher costs for BMI and WHR one unit increases in our IV model than in the OLS model. Because the association of genetic variants with BMI and WRH is still weak, results have to interpreted carefully.

## Acknowledgements

We thank Konstantin Strauch, Thomas Meitinger, Harald Grallert, Christine Meisinger, and Annette Peters for provding the data.

